# Metabolomic profile for understanding heart failure classifications

**DOI:** 10.1101/2024.11.13.623517

**Authors:** Maria V. Kozhevnikova, Yuri N. Belenkov, Ksenia M. Shestakova, Anton A. Ageev, Pavel A. Markin, Anastasiia V. Kakotkina, Ekaterina O. Korobkova, Natalia E. Moskaleva, Ivan V. Kuznetsov, Natalia V. Khabarova, Alexey V. Kukharenko, Svetlana A. Appolonova

## Abstract

**Background:** The existing classifications of heart failure (HF) remain a topic of debate within the cardiology community. Metabolomic profiling (MP) offers a means to define HF phenotypes based on pathophysiological changes, allowing for more precise characterization of patient groups with similar clinical profiles. MP may thus aid in refining HF classifications and offer a novel approach to phenotyping.

**Methods:** MP was performed to 408 patients with different stages of HF. Patients with symptomatic HF were divided into phenotypes by left ventricle ejection fraction (LVEF). Liquid chromatography combined with mass-spectrometry were used for the MP. Data were analyzed using machine learning. The relationship between the incidence of all-cause death and LVEF trajectory changes and metabolomic clusters was evaluated. Follow-up period was 542 days [16;1271] in average.

**Results:** The classification model achieved an AUC ROC - 0.91 for distinguishing of Stage A from Stage B and an AUC ROC - 0,97 for Stage B vs. Stage C using metabolomic analysis, model’s performance for differentiating Stages C and D was lower (AUC ROC 0.81). For HF phenotypes, the HFrEF, HFmrEF, and HFpEF model demonstrated moderate accuracy (AUC ROC 0.74), whereas the model distinguishing HFpEF from HF with EF <50% showed good precision. The HFrEF vs. HF with EF >40% model, however, displayed low accuracy. Biostatistical processing of MP identified four metabolomic clusters, and 26 metabolites demonstrated the greatest significance (metabolites of the kynurenine and serotonin pathways of tryptophan catabolism, glutamine, riboflavin, norepinephrine, serine, long- and medium-chain acylcarnitines). Patients with reduced LVEF had the poorest prognosis (HR 1,896; 0,711–5,059), with an LVEF decrease linked to a threefold rise in all-cause mortality risk. Cluster 3 was associated with a 2,880-fold increase in all-cause mortality.

**Conclusions:** Our findings suggest that MP provides an effective alternative approach for stratifying HF patients by stage. The observed metabolic similarities between HFpEF and HFrEF phenotypes highlight limitations in the current classification, underscoring the need to refine HF phenotyping into two primary categories. Hierarchical clustering by metabolomic profile produced a high-accuracy model, supporting MP as a valuable tool for HF classification.

**Novelty and Significance:** *What Is Known?:*

- Classifying HF by stages offers the advantage of incorporating preventive aspects, however, diagnosing Stage B can be challenging due to the necessity of analyzing numerous parameters for verification.
- The classification by LVEF, particularly distinguishing HFmrEF, remains controversial because of limited evidence for specific treatment strategies.
- Metabolomic profiling (MP) offers a means to identify unique pathophysiological changes across phenotypes, presenting a promising approach for diagnosing HF and developing targeted therapies.

*What New Information Does This Article Contribute?:*

- This study demonstrates high accuracy in HF stage classification by integrating chromatography-mass spectrometry data with bioinformatic analysis through multiparametric machine learning (ML) models.
- Similarities between metabolomic profiles of patients with LVEF <40% and those with LVEF 41–49% suggest pathophysiological overlap between HFmrEF and HFrEF.
- MP, enhanced by ML, allows precise differentiation between HFpEF and patients with EF <50%, with an AUC ROC of 0.96.
- Hierarchical clustering based on metabolomic profiles identified four distinct HF phenotypes, reflecting unique pathophysiological pathways with high accuracy (AUC ROC 0.96). This work highlights how metabolomic analysis provides insight into HF’s biochemical landscape, aiding classification and offering potential new pathways for therapeutic intervention. The findings underscore MP’s potential to improve HF phenotyping and understanding of disease mechanisms.

**Graphical abstract.**
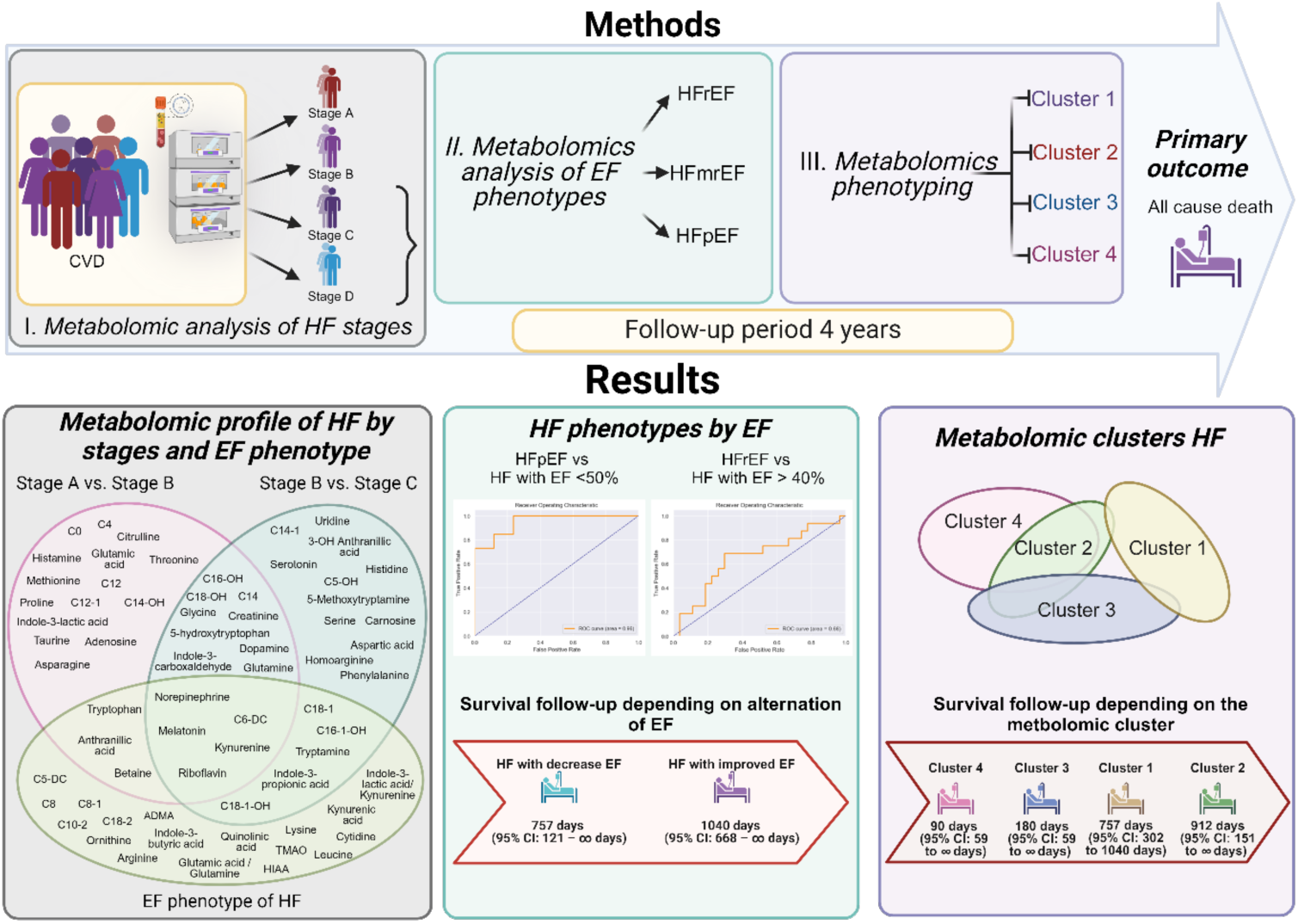
EF – ejection fraction; HF – heart failure; HFmrEF - heart failure with mid-range ejection fraction; HFpEF - heart failure with preserved ejection fraction; HFrEF - heart failure with reduced ejection fraction.

## Introduction

Heart failure (HF) is a complex heterogeneous syndrome, which is determined by a wide variety of etiological factors, variants of myocardial remodeling, activation of various regulatory systems. The course of the disease, prognosis and treatment options are determined by the specific variant of HF. For this reason, special attention is paid for the classification of HF. In order to standardize classification approaches, the regulatory document was issued in 2021, according to which HF can be classified by stages, left ventricle ejection fraction (LVEF), and functional class.^1^ The effectiveness of the renin-angeotensin-aldosterone system inhibitors and implantable devices in patients with HF with reduced EF (HFrEF) and the lack of improvement in prognosis in patients with mildly reduced and preserved EF causes the separation of patients by LVEF. However, with the advent of new methods of treatment and new medicines that are effective in the cross all spectrum of EF the classification of LV began to lose its relevance. In addition, modern therapy can significantly improve LVEF, which is the reason for the transition of patients from one phenotype to another. In this regard, the AHA/ACC/HFSA decided to include HF with improved EF to the classification as a separate phenotype.^2^

Thus, existing classifications have a number of shortcomings and do not provide a complete understanding of the characteristics of pathophysiological changes in various conditions. Perhaps, for the further development of therapeutic approaches, a new look at the classification of HF with a shift in emphasis towards molecular mechanisms is needed. A deeper understanding of the pathophysiological features of various HF phenotypes by EF and stages can be achieved using omics technologies. Genomics, transcriptomics and metabolomics provide biological interpretation of changes in tissues, organs and systems. Metabolomic profiling of plasma is an accessible method for assessing metabolic disorders at the systemic level and provides information about the pathophysiological changes that occur during the disease. A deeper understanding of the pathophysiological features of various HF phenotypes by EF and stages can be achieved using omics technologies.

In this regard, the purpose of this study was to analyze changes in metabolic pathways in various classifications of HF, to improve understanding of the heterogeneous nature of HF and search for potential therapeutic targets.

## Methods

### Study population

We included 408 patients with different stages of HF according to the American Heart Association/American Heart Failure Association classification.^2^ Stage A was defined by absence of HF symptoms, structural heart disease, or cardiac biomarkers increase. Stage B was detected by absence of symptoms or signs of HF but presence of structural changes: LAVI ≥29 mL/m^2^, LVMI >115/95 g/m^2^, relative wall thickness (RWT) >0.42, LV wall thickness ≥12 mm, average E/e′ ≥15 for increased filling pressures, septal e’ <7 cm/s, lateral e’ <10 cm/s, TR velocity >2.8 m/sec, pulmonary artery systolic pressure (PASP) >35 mmHg or brain natriuretic peptide (BNP) ≥35 pg/mL, NT-proBNP ≥125 pg/mL. Stage C characterized by structural heart disease with current or previous symptoms of HF. Stade D was defined by marked HF symptoms that interfere with daily life and with recurrent hospitalizations despite attempts to optimize guideline-directed medical therapy.^2^ Patients with presence of signs of lung congestion or peripheral oedema, hepatomegaly, jugular venous distention were classified by EF. HFrEF was defined by a decrease of LVEF <40%. HF with mid-range EF (HFmrEF) was determined in LVEF 40-49%. HF with preserved EF (HFpEF) was verified in LVEF ≥50% and N-terminal pro-B-type natriuretic peptide (NT-proBNP) >125 pg/mL or at least one of the additional items: enlargement of the left atrium (LA) (LA volume index (LAVI) >34 ml/m^2^), hypertrophy of the LV (LV mass index (LVMI) >115 in men and >95 g/m^2^ in women), early diastolic velocity of movement of the fibrous ring of the mitral valve (e’) at the level of the interventricular septum (<7 cm/s) and the posterior wall (<10 cm/s), average E/e’ (≥15), maximum speed of tricuspid regurgitation (TR) velocity (>2.8 m/s).^3^ Hypertension and coronary artery disease (CAD) was defined according to the guidelines.^4,5^

Exclusion criterion were: acute forms of CAD, cardiomyopathies; valvular heart disease; myocarditis, pericarditis; stroke within the previous 6 months; acute renal failure, chronic kidney disease stage 5; chronic pulmonary heart disease; hepatic cellular failure; bronchial asthma, chronic obstructive pulmonary disease; gastric ulcer in the acute phase; chronic pancreatitis in the acute phase; cancer; type 1 diabetes mellitus; thrombocytopenia of any origin, hemorrhagic syndrome; chronic viral and bacterial infections; autoimmune diseases; mental illness.

The study was conducted with the Declaration of Helsinki of the World Medical Association principles, adopted at the 18th General Assembly of the WMA, was approved by the Local Ethics Committee of Sechenov University protocol №34-20 (09.12.2020) and was carried out by decision of the Academic Council of Sechenov University. All study participants gave written informed consent.

### Metabolomic analysis

Quantitative analysis of metabolites was performed by target chromato-mass-spectrometric method according to with some modifications.^6^

10 μL of the blood plasma sample (or calibration standards and quality control (QC) samples) and 40 μL of ISTD mix in methanol were pipetted in Eppendorf tubes. After drying the samples in vacuum concentrator 50 μL of PITC derivatization solution (5% phenyl isothiocyanate in the mixture of acetonitrile, water and pyridine, 1:1:1) was added and kept at room temperature for 20 min. After another 1-h drying in vacuum concentrator 100 μL of methanol with 5mM ammonium acetate and 100 μL of water were added to each sample. After 30 min extraction samples were centrifuged at 13000 rpm for 10 min and supernatant was analyzed by liquid chromatography mass spectrometry (LC-MS/MS).

An Agilent 1200 series HPLC system and Agilent 6460 triple-quadruple mass-spectrometer (Palo Alto, CA, U.S.A.) were used for LC-MS/MS analysis. Acquity UPLC BEH C18 2,1 × 50 mm; 1,7 mcm (Waters Corp., U.S.A) column with Acquity UPLC BEH C18 (2,1 × 5 mm; 1,7 mcm) precolumn was used for chromatographic separation. The UPLC parameters were as follows: solvent A 0.1% (v/v) formic acid in water, and solvent B 0.1% (v/v) formic acid in acetonitrile. The column temperature was maintained at 40°C. The gradient program was as follows: 0.5 min - 1% B, 1min – 30 % B, 3 min - 75% B, 3,5 min - 95% B, 4.5 min - 95% B, 4,6 min - 1% B, 6 min - 1% B. Flow rate was 500 μL/min, and the sample injection volume was 10 μL.

A mass spectrometric detector with a triple quadrupole in the scheduled MRM mode was used for analysis (Table S1). For Agilent Jet stream ion source the capillary voltage was 3500V and nebulizer pressure was 30 psi. The Gas Temp, Gas Flow, Sheath Gas Heater and Sheath Gas Flow were 300°C, 11 l/min, 300°C and 11 l/min, respectively. The fragmentor voltage, collision energy, MRM precursor ion (Q1), and fragment ion (Q3) were optimized and set individually for each analyte and isotope-labeled ISTD.

Quantitative analyses were done in MassHunter software (Agilent, Palo Alto, CA, U.S.A.) based on the peak area ratios of the targeted analyte compared to its isotope-labeled ISTD. Calibration regression was built for each analyte within its plasma concentration range. For acylcarnitine analysis quantification was done by semiquantitative method based on the peak area ratios of the target analyte and its isotope-labeled standards only without calibration curves.

Methods were validated for selectivity, linearity, precision, accuracy, recovery, matrix effect and stability according to US FDA and EMA guidelines for bioanalytical method validation (EMA, 2019; USFDA, 2018). Detection limit and linear range are selected for each compound according to physiological plasma concentrations.

We included 408 patients with different stages of HF according to the American Heart Association/American Heart Failure Association classification.^2^ Stage A was defined by absence of HF symptoms, structural heart disease, or cardiac biomarkers increase. Stage B was detected by absence of symptoms or signs of HF but presence of structural changes: LAVI ≥29 mL/m^2^, LVMI >115/95 g/m^2^, relative wall thickness (RWT) >0.42, LV wall thickness ≥12 mm, average E/e′ ≥15 for increased filling pressures, septal e’ <7 cm/s, lateral e’ <10 cm/s, TR velocity >2.8 m/sec, pulmonary artery systolic pressure (PASP) >35 mmHg or brain natriuretic peptide (BNP) ≥35 pg/mL, NT-proBNP ≥125 pg/mL. Stage C characterized by structural heart disease with current or previous symptoms of HF. Stade D was defined by marked HF symptoms that interfere with daily life and with recurrent hospitalizations despite attempts to optimize guideline-directed medical therapy.^2^ Patients with presence of signs of lung congestion or peripheral oedema, hepatomegaly, jugular venous distention were classified by EF. HFrEF was defined by a decrease of LVEF <40%. HF with mid-range EF (HFmrEF) was determined in LVEF 40-49%. HF with preserved EF (HFpEF) was verified in LVEF ≥50% and N-terminal pro-B-type natriuretic peptide (NT-proBNP) >125 pg/mL or at least one of the additional items: enlargement of the left atrium (LA) (LA volume index (LAVI) >34 ml/m^2^), hypertrophy of the LV (LV mass index (LVMI) >115 in men and >95 g/m^2^ in women), early diastolic velocity of movement of the fibrous ring of the mitral valve (e’) at the level of the interventricular septum (<7 cm/s) and the posterior wall (<10 cm/s), average E/e’ (≥15), maximum speed of tricuspid regurgitation (TR) velocity (>2.8 m/s).^3^ Hypertension and coronary artery disease (CAD) was defined according to the guidelines.^4,5^

Exclusion criterion were: acute forms of CAD, cardiomyopathies; valvular heart disease; myocarditis, pericarditis; stroke within the previous 6 months; acute renal failure, chronic kidney disease stage 5; chronic pulmonary heart disease; hepatic cellular failure; bronchial asthma, chronic obstructive pulmonary disease; gastric ulcer in the acute phase; chronic pancreatitis in the acute phase; cancer; type 1 diabetes mellitus; thrombocytopenia of any origin, hemorrhagic syndrome; chronic viral and bacterial infections; autoimmune diseases; mental illness.

### Clinical follow-up

Patients with symptomatic HF (n=218) had a follow-up period. The primary outcomes were all cause death. The average follow-up period was 542.37 [16;1271] days. When multiple events occurred, patients were censored at the time of the first event. Follow-up events were adjudicated by an independent trained investigator.

### Statistical analysis

Quantitative indicators were assessed for compliance with normal distribution using the Shapiro-Wilk test (if the number of subjects was less than 50). Comparison of two groups according to a quantitative indicator with a normal distribution, provided that the variances were equal, was performed using the Student t-test, and when the variances were unequal, there was applied the Welch t-test. Comparison of two groups for quantitative indicators whose distribution differed from normal was performed using the Mann-Whitney U test. Comparison of several groups on a quantitative indicator with a normal distribution was performed using one-way analysis of variance, post-hoc comparisons were carried out using Fisher’s test (assuming equal variances) and Welch’s test (if unequal variances). To compare quantitative indicators of several groups, the distribution of which differed from normal, the nonparametric Kruskal-Wallis test was used. If statistically significant differences were identified, further post hoc pairwise comparison of groups was carried out using Dunn’s test with Holm’s correction. Pearson chi-square was used to compare categorical variables across multiple independent groups. Differences were considered significant at p<0.05. The direction and strength of the correlation between two quantitative indicators were assessed using the Pearson correlation coefficient (if the compared indicators were normally distributed) or the Spearman correlation coefficient (if the compared indicators were not normally distributed). The connection more than 0.3 were taking into account. Patients’ survival function was assessed using the Kaplan-Meier method. The analysis of patient survival was carried out using the Cox regression method, which involves predicting the risk of an event for the object in question and assessing the influence of predetermined independent variables (predictors) on this risk. Risk is viewed as a time-dependent function. Missing data were not included in the models. Data were analyzed with SPSS version 25.0 (IBM Corp., Armonk, NY, USA) and R 3.6.2 (R Foundation for Statistical Computing, Vienna, Austria).

Raw LC-MS/MS data preprocessing included normalization and replacement of outliers and missing values via pandas (v.2.2.3), numpy (v.2.1.2), scipy (v.1.14.1) packages in python (v.3.11). QC samples were analyzed each ten samples to control stability of the instrument and to normalize the variations in analyzed batch.

Further principal component analysis was applied to explore trends in the results of metabolomic profiling in the desired groups of patients. Additionally, to assess the diagnostic accuracy of different classifications of HF stages ML models based on the random forest algorithms were built. For this reason, the desired datasets were randomly divided into discovery (70%) and test datasets (30%). Hyperparameters for the random forest models was tuned using cross-validated (k=30) GridSearchCV function.

The random forest algorithm is a nonparametric ensemble method is based on a combination of several decision trees, called “random trees”. Each tree is built based on a random subsample of the training data and a random subset of features, that implements “randomness” into the learning of each tree. It serves for the reduction of the correlation between trees and increases the diversity of models.

The assessment of classification quality was performed using common quality metrics, including confusion matrix, area under the receiver operating characteristics curve (AUC ROC), accuracy, f1-score and recall. The principal components analysis (PCA), random forest models were performed using the scikit-learn package (v.1.5.2) in Python (v.3.11).

Hierarchal cluster analysis (HCA) is a clustering method that explores the organization of samples into groups therefore providing a hierarchy usually represented as a dendrogram. Here, the HCA was performed using the Ward minimum variance method and preliminary reduction of the data dimensionality to nine principal components via Simca-P+ software (version 14.1, Umetrics).

## Results

### Study population

Demographic and clinical characteristics of all subjects are shown in Table 1. Stage A was presented predominantly by hypertensive patients and coronary artery disease (CAD) patients (130 patients, 31.9%). The stage B was represented predominantly by patients with CAD and to a lesser extent by patients with hypertension (60 patients, 14.7%). HFrEF had 81 patients (37.0%), HFpEF was detected in 87 patients (40.0%), HFmrEF was detected in 50 patients (23%).

**Table 1.** Population characteristics. ACEi, angiotensin-converting enzyme inhibitor; ARB, angiotensin receptor blocker; ARNI, angiotensin receptor–neprilysin inhibitor; BMI, body mass index; CAD, coronary artery disease; CABG – coronary artery bypass grafting; CCB, calcium channel blocker; eGFR, estimated glomerular filtration rate; HDL, high-density lipoprotein; HFmrEF - heart failure with mid-range ejection fraction; HFpEF, heart failure with preserved ejection fraction; HFrEF, heart failure with reduced ejection fraction; CRP, C-reactive protein; LDL, low-density lipoprotein; MI, myocardial infarction; MRA, mineralocorticoid receptor antagonist; NT- proBNP, N-terminal pro-B-type natriuretic peptide; PCI, percutaneous coronary intervention; SGLT2i, sodium/glucose cotransporter-2 inhibitors; TIA, transient ischemic attack; WBC, white blood counts. • * P < 0.01 vs. Stage A. • ** P < 0.01 vs. Stage B. • *** P<0.01 vs. Stage C.

| Variable | Stage A<br>(n=130) | Stage B<br>(n=60) | Stage C<br>(n = 118) | Stage D<br>(n = 100) | P-value |
| --- | --- | --- | --- | --- | --- |
| **Demographics** |  |  |  |  |  |
| Age, years | 62 [53-68] | 67 [65–74] * | 68 [61–72]* | 69 [65–73]* | **<0.00000**<br>**1** |
| Male sex, n (%) | 60 (46.2) | 32 (53.3) | 70 (59.3) | 62 (62.0) | 0.07 |
| BMI, kg/m<sup>2</sup> | 30 [26-32] | 32 [28–25] | 32 [27–36]* | 31 [28–35] | **<0.009** |
| Smoking, n (%) | 30 (25.2) | 12 (20.0) | 25 (21.4) | 12 (12.0) | 0.10 |
| **Clinical evaluation** |  |  |  |  |  |
| Obesity | 58 (44.6) | 40 (66.7) * | 67 (56.8) | 53 (53.0) | **0.03** |
| Arterial hypertension | 99 (76.2) | 50 (83.3) | 117 (99.1) | 100 (100) | **<0.0001** |
| Dyslipidaemia<sup>a</sup> | 99 (76.1) | 49 (83.1) * | 100 (84.7)<br>* | 71 (71.0) *** | **<0.00000**<br>**1** |
| Impaired glucose tolerance | 14 (11.9) | 11 (18.3) | 20 (16.9) | 21 (21.0) |  |
| Diabetes mellitus | 25 (21.2) | 17 (28.3) | 43 (36.4) * | 49 (49.0) *** | **<0.00002** |
| Stroke/TIA | 1 (2.1) | 8 (18.6) | 20 (16.9) | 16 (16.0) | 0.06 |
| CAD | 34 (26.2) | 48 (80.0) | 50 (83.3) | 100 (100.0) | **<0.0001** |
| PCI | 12 (9.2) | 22 (36.7) * | 19 (16.1) *<br>* | 27 (27.0) * | **<0.00002** |
| CABG | 2 (4.4) | 3 (6.4) | 8 (10.3) | 5 (5.0) | 0.49 |
| Previous MI | 10 (10.4) | 26 (46.4) * | 48 (61.5) * | 77 (77.0) *** | **<0.00000**<br>**1** |
| HFpEF | 0 | 0 | 69 (58.5) | 18 (18.0) | **<0.00000**<br>**1** |
| HFmrEF | 0 | 0 | 21 (17.8) | 29 (29.0) |  |
| HFrEF | 0 | 0 | 28 (23.7) | 53 (53.0) |  |
| Blood tests |  |  |  |  |  |
| Haemoglobin, g/L | 141.0<br>[132.0–<br>153.0] | 144.0<br>[133.0–<br>154.0] | 138<br>[129–<br>148.0] | 132<br>[119.75–<br>144.0]<br>***** | **0.000006** |
| WBC, *10<sup>9</sup> | 6.2<br>[5.4–7.53] | 7.2<br>[5.97–8.81]<br>* | 7.8<br>[6.4–9.17]* | 8.0<br>[6.8–<br>9.65]*,*** | **<0.00001** |
| K<sup>+</sup>, mMol/L | 4.6<br>[4.38-4.90] | 4.5<br>[4.30-4.90] | 4.5<br>[4.30-4.97] | 4.4<br>[4.0-<br>4.7]*,*** | **0.00058** |
| Total cholesterol,<br>mmol/L | 5.27± 1.28 | 4.57 ± 1.36* | 4.35 ± 1.37<br>* | 3.64 ± 1.08**<br>* *** | **<0.00000**<br>**1** |
| LDL, mg/dL | 3.25<br>[2.44-3.75] | 2.48<br>[1.88-3.27]* | 2.94<br>[1.9-3.59]*** | 2.13<br>[1.52-2.93]* | **<0.0001** |
| HDL, mg/dL | 1.42<br>[1.18-1.67] | 1.17<br>[1.01-1.55] | 1.04<br>[0.86-1.30]* | 0.95<br>[0.72-1.23]* | **<0.00000**<br>**1** |
| Triglycerides,<br>mg/dL | 1.3<br>[1.02-1.91] | 1.23<br>[1.00-1.83] | 1.36<br>[0.98-1.95]* | 1.09<br>[0.82-1.40]* | **<0.0053** |
| Creatinine,<br>mkmol/L | 91.95<br>[81.42-101.6] | 97.10<br>[84.48–<br>110.72] | 99.5<br>[88.25–<br>114.00] | 111.0<br>[93.75-127.25]<br>***,*** | **<0.00000**<br>**1** |
| eGFR,<br>mL/min/1.73 m<sup>2</sup> | 69.37± 15.0<br>9 | 62.42± 19.28<br>* | 60.76± 15.7<br>1 | 54.02± 15.82<br>*,**,*** | **<0.00000**<br>**1** |
| Fasting blood<br>glucose, mmol/L | 5.6 [5.2-6.3] | 5.8 [5.1–<br>6.52]* | 6.0 [5.3–<br>7.2] | 6.0 [5.0–8.2] | 0.165 |
| Uric acid,<br>mkmol/L | 328.5<br>[274.8-385.2] | 353.8<br>[311.2–<br>406.9] | 410.0<br>[311.2–<br>482.0]* | 456.0<br>[361.0–<br>529.0]*,***,**<br>* | **<0.00000**<br>**1** |
| CRP, mg/L | 1.57 [0.66-4.00] | 1.6 [0.70–<br>3.00] | 4.00 [1.50–<br>7.10]* | 9.13 [4.49–<br>22.54]*,***<br>*** | **<0.00000**<br>**1** |
| Ferrum, mkmol/L | 17.55<br>[15.30-22.50] | 14.10<br>[11.70-22.20] | 14.00 [9.7-20.95]* | 6.50 [4.00-14.70]*,***,*<br>** | **0.000014** |
| NT-proBNP,<br>pg/mL | 51 [48-56] | 101 [46–<br>143] | 1300 [376–<br>3953]*,*** | 3733 [1341–<br>7250]*,***,**<br>* | **<0.00000**<br>**1** |
| **Therapy** |  |  |  |  |  |
| Beta-blocker | 63 (52.5) | 44 (73.3)* | 101 (85.6)* | 87 (87.0)* | **<0.00000**<br>**1** |
| ACEi | 67 (56.3) | 34 (56.7) | 50 (42.4) | 30 (30.0) | **0.00036** |
| ARB | 31 (25.8) | 14 (23.3) | 37 (31.4) | 18 (18.0) | 0.153 |
| ARNI | 0 (0.0) | 0 (0.0) | 27 (34.6) | 49 (49.0) | **<0.0001** |
| MRA | 5 (5.0) | 15 (25.0) | 77 (65.3) | 83 (83.0) | **<0.0001** |
| SGLT2i | 1 (2.1) | 7 (16.3) | 18 (15.3) | 19 (19.0) | 0.051 |
| Digoxin | 0 | 0 | 16 (13.6) | 31 (31.0) | **0.01** |
| CCB | 32 (26.7) | 20 (33.3) | 38 (32.2) | 12 (12.0)<br>*,**,*** | **0.0025** |
| Oral anticoagulant | 24 (20.0) | 26 (43.3)* | 75 (63.6)* | 78 (78.0)<br>*,**,*** | **<0.0001** |
| Statins | 55 (45.8) | 17 (28.3) | 45 (38.1) | 52 (52.0)** | **0.0178** |
| Furosemide | 0 | 0 | 78 (66.1) | 85 (85.0) | **<0.0001** |
| Amiodaron | 11 (9.2) | 8 (13.3) | 14 (11.9) | 2 (2.0)***,*** | **0.033** |

Values are mean ± standard deviation, *n* (%), or median [25th quartile–75th quartile].

### Metabolomic analysis design and data preparation

To implement the objectives of the study, a metabolomic analysis design was constructed, according to which the creation of groups for comparison was determined by a specific task at each stage. At the first stage, the comparing analysis of HF stages was performed. The second stage of the analysis included only patients of the HF C and D groups, who were divided into 3 groups according to the LVEF phenotype. And, finally, at stage 3, patients with HF stage C and HF stage D were clustered according to the metabolomic profile (Figure 1).

**Figure 1.**
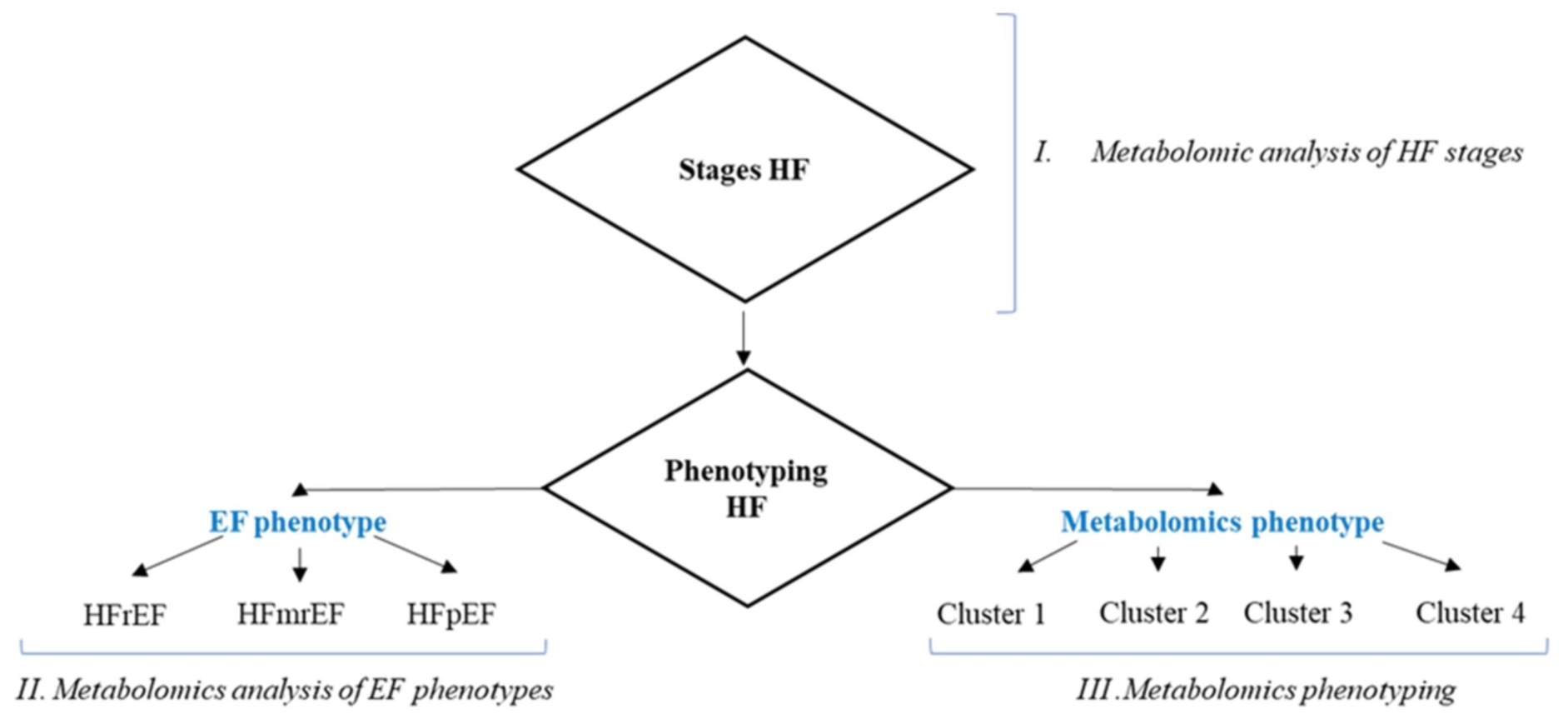
Design of the metabolomic analysis. EF - ejection fraction; HF - heart failure; HFmrEF - heart failure with mid-range ejection fraction; HFpEF - heart failure with preserved ejection fraction; HFrEF - heart failure with reduced ejection fraction.

### Metabolomic profile in patients with different stages of HF

The conducted principal component analysis showed weak differences in the four desired groups of patients (Figure S1). Further we have learned the machine learning (ML) classification model to assess the possibility of separation HF patients in according to the provided methodology. The utilized ML algorithm represented random forest classifier with the hyperparameters and quality control metrics presented in the Table S2 and S3, respectively. Through the following deeper investigation of the data, we also assessed the diagnostic power between patients from Stage A vs. Stage B groups (Figure 2A), Stage B vs. Stage C groups (Figure 2B), and Stage C vs. Stage D groups (Figure 2C). The calculated AUC ROC metrics showed weak diagnostic power for separating patients between stage C and stage D. Data from the most significant metabolites in all models are reported in Table S4-S6. This result demonstrated that classification models of ML can accurately predict the presence of stage B od stage C.

**Figure 2.**
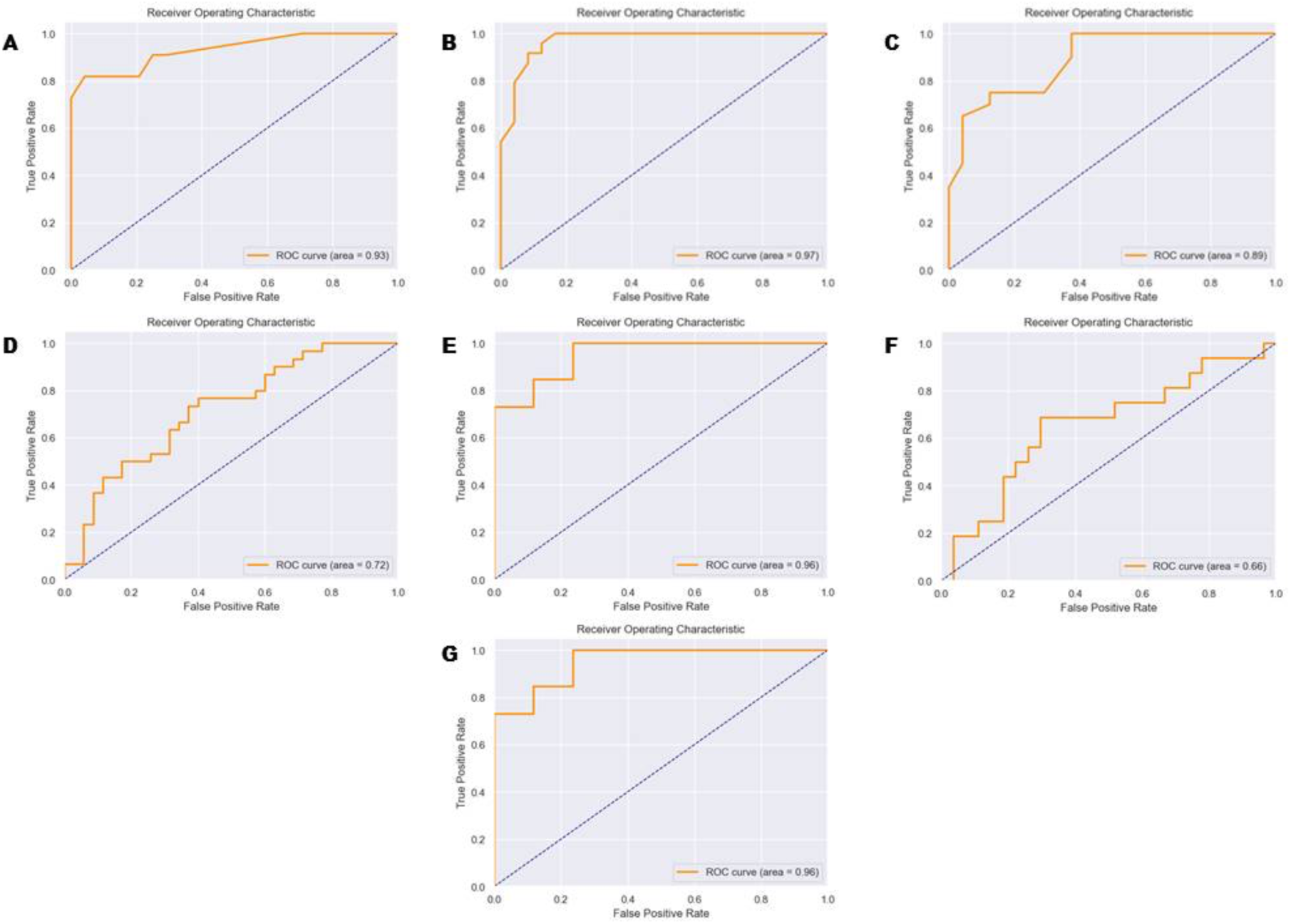
ROC curve of HF in the studied subgroups. **A.** Classification model – Stage A vs. Stage B groups. Confusion matrix: Accuracy 0.88, Recall 0.88, AUC ROC 0.93, F1 0.88. **B.** Classification model – Stage B vs. Stage C groups. Accuracy 0.89, Recall 0.89, AUC ROC 0.97, F1 0.89. **C.** Classification model – Stage C vs. Stage D groups. Confusion matrix: Accuracy 0.79, Recall 0.79, AUC ROC 0.89, F1 0.79. **D.** Classification model – HFpEF vs. HFmrEF vs. HFrEF. Confusion matrix: Accuracy 0.66, Recall 0.66, AUC ROC 0.72, F1 0.64. **E.** Classification model - HFpEF vs. HF with EF <50%. Confusion matrix: Accuracy 0.88, Recall 0.88, AUC ROC 0.96, F1 0.88. **F.** Classification model - HFrEF vs. HF with EF > 40%. Confusion matrix: Accuracy 0.70, Recall 0.70, AUC ROC 0.66, F1 0.68. **G.** Classification model – Cluster 1 vs. Cluster 2 vs. Cluster 3 vs. Cluster 4. Confusion matrix: Accuracy 0.95, Recall 0.95, AUC ROC 0.96, F1 0.95.

### Metabolomic profile in patients with different EF phenotypes

For better classification of patients from stage C and stage D using results of the targeted metabolomic profiling we applied the alternative classification method specified only for separation of patients with HF. Thus, patients from stage C and stage D groups were separated into three phenotypic classes in accordance to the EF and labeled as HFpEF, HFmrEF and HFrEF.^3^ The assessment of the possible application of such classification was performed using the ML modelling as is represented above. Diagnostic quality of the multiclass model is presented in Figure 2D. The quality metrics showed weak diagnostic accuracy of the model thus we made a hypothesis that metabolomic profile in HFmrEF can be similar to HFpEF or HFrEF types. So, two classification models were found: HFpEF vs. HF with EF <50% and HFrEF vs. HF with EF > 40%. The first model (HFpEF vs. HF with EF <50%) had good precision (Figure 2E), while the model HFrEF vs. HF with EF > 40% had poor accuracy (Figure 2F). This results suggested support the model HFpEF vs. HF with EF <50%. The key metabolites forming the classification model of HFpEF vs. HF with EF < 50% presented in the Table S7.

### Clustering by metabolomic profile

Alternatively, we applied the cluster analysis method to identify new classification based on the results of the plasma metabolomic profiling. In this case, we performed hierarchical clustering by metabolomic profile based on the first main component that divided all patients with symptomatic HF (stage C and D) into four clusters (Figure 3A, 3B). The new classification model had AUC ROC 0.96 with metrics: max_depth=30, max_features=20, min_samples_leaf=1, n_estimators=20, random_state=42. (Figure 2G). Further, the 26 most significant metabolites for this classification were identified, belonging mainly to the class of acylcarnitine’s, metabolites of the tryptophan-kynurenine and tryptophan-serotonin pathways, and several amino acids (Table S8).

**Figure 3.**
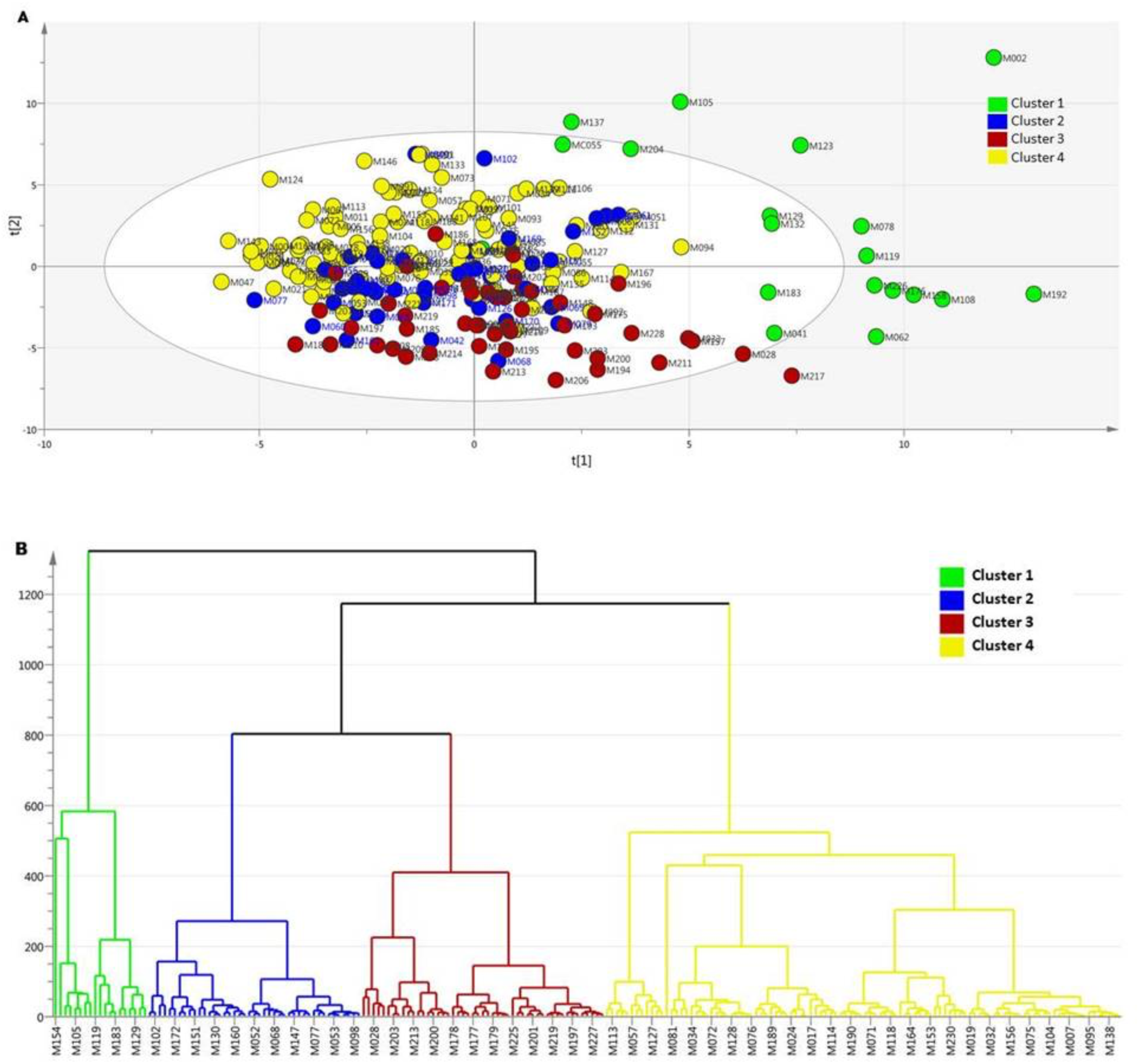
Clustering of patients with heart failure by metabolomic profile. **A.** PCA by metabolomic profile. **B.** Hierarchical clustering by metabolomic profile. PCA - principal components analysis.

Thus, it was possible to divide all patients with HF into 4 clusters using biostatistical processing of the metabolomic profile. In this classification, 26 metabolites demonstrated the greatest significance. Further, to assess the nature of metabolic disorders characteristic of each cluster, a comparative analysis of 4 clusters for significant metabolites was carried out. Metabolites of the kynurenin pathway of tryptophan catabolism (3-OH-anthranilic acid, quinolinic acid, and xanthurenic acid) were significantly increased in cluster 3 compared to other clusters. At the same time, serotonin pathway metabolites (5-hydrocytryptophan, 5- methoxytryptamine) were significantly reduced in cluster 1 compared to the other groups. Glutamine was significantly reduced in the second group and the third clusters compared to clusters 1 and 4. An increase in riboflavin was characteristic of cluster 3. Norepinephrine was statistically significantly elevated in cluster 2 compared to all groups studied, while cluster 3 had the lowest values. Cluster 3 was characterized by a decrease in serine compared to other subgroups. Finally, long- and medium-chain acylcarnitine, as well as the isovalerylcarnitine, tiglylcarnitine, and glutarylcarnitine metabolites, were statistically significantly higher in cluster 4 compared to the other clusters (Figure 4, Table S9).

**Figure 4.**
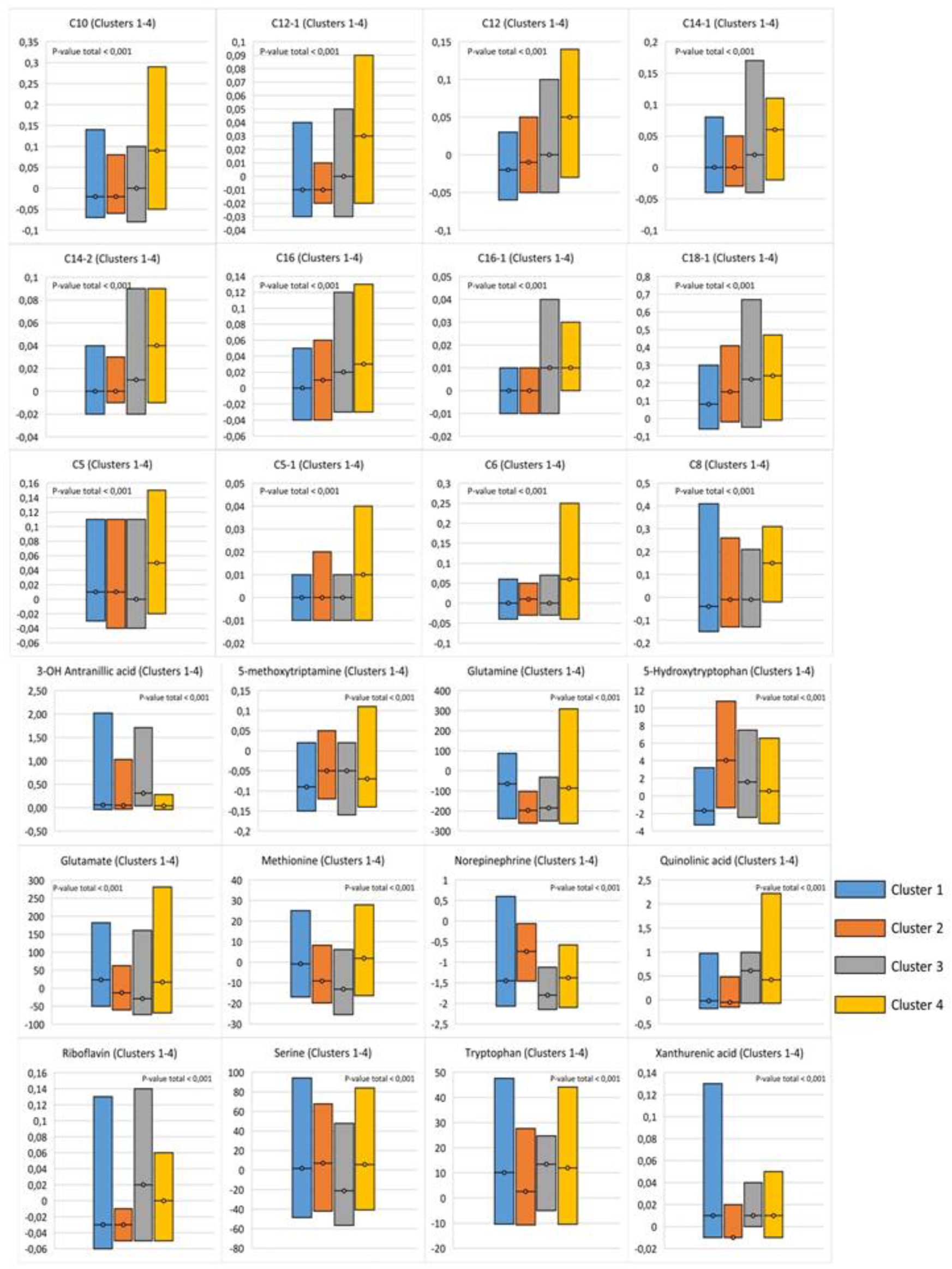
Comparative analysis of significant metabolites in the classification model of clustering.

### Survival of patients with chronic heart failure

During a median follow-up of 542.37 [16; 1271] days, 57 (26.14%) patients had outcomes event (all causes death), the annual mortality rate was 14% (n=31). For the Kaplan-Meier assay, patients who discontinued follow-up within 1 month of hospital discharge were excluded from the analysis (n=24). The 25th percent was 481.64 days. The analysis showed that the overall survival rate at 2 years was 74.8%. At the same time, half of the deaths occurred in the first 2 months of follow-up (n=29). The three-year risk of death in patients with HF was 41.9%.

The study design involved only telephone contact with the patient at the end of follow-up to assess outcomes development. However, information from the medical records of patients from the HF group who regularly sought medical care at the clinical sites of the study during the follow-up period was analyzed (n = 98). It was found that 49 (50%) patients did not show significant deviations in LVEF levels during follow-up (less than 5%), 16 (16.3%) patients had a decrease in EF, and 33 (33.7%) patients showed improvement or recovery of LVEF during treatment. In the analysis of the therapy taken, it was found that 8 patients (8.2%) patients regularly use the GDMT, 31 (31.6%) of the patient used only two classes of recommended therapy, the remaining 59 (60.2%) patients used 3 of the four recommended classes of drugs. GDMT treatment group had a better survival rate of 87.5% compared to patients taking 3 classes of drugs (54.9%) and patients taking 2 classes of drugs (52.0%). During follow-up, myocardial infarction developed in 8 (8.2%) patients, 7 of whom underwent percutaneous coronary intervention and stenting of the infarction-binding artery. Two patients (2%) underwent coronary artery bypass graft as planned. There was no significant association of EF changes with therapy, MI or revascularization.

To assess the course of HF in the studied subgroups (EF persistent, EF decreased, EF improved), the Kaplan-Meier analysis was carried out (Figure 5A). The analysis showed that within 1000 days, patients with decreased LVEF had the worst prognosis, regardless of the baseline level of EF. However, survival was later comparable between patients with reduced and improved EF (Figure 5A).

**Figure 5.**
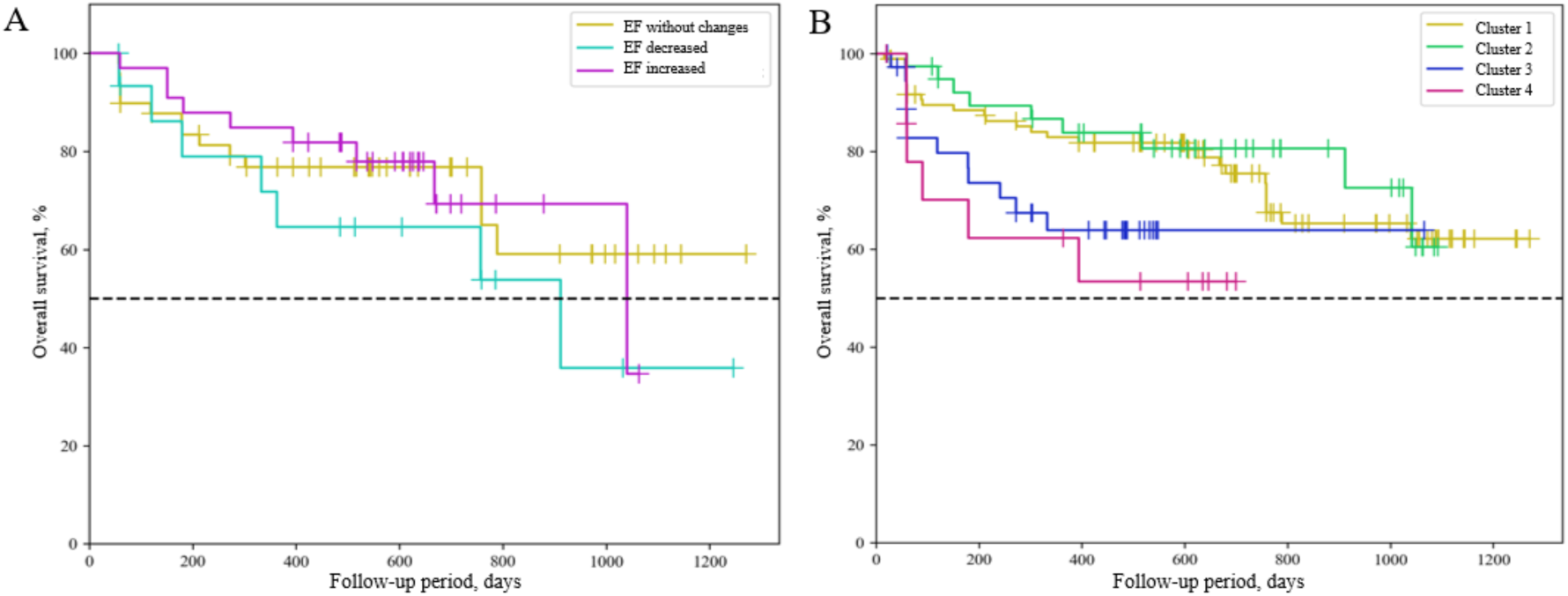
Overall survival analysis of HF in the studied subgroups. Kaplan-Meier curves of HF by EF persistent, EF decreased and EF improved (A) and by metabolomic cluster (B) subgroups. EF – ejection fraction, HF - heart failure.

Patients with persistent EF had the most favorable prognosis during the follow-up period. However, the differences in overall survival assessed by the likelihood ratio test were not statistically significant (p = 0.337096). The analysis showed an increased risk of all-cause death in patients with reduced LVEF (HR 1.896; 0.711 to 5.059) compared to patients with improved EF (HR 0.829; 0.325 to 2.114). At the same time, there was no statistically significant difference between the groups. However, the lack of differences may be due to the small size of the groups.

The analysis showed that metabolomic cluster 4 was associated with the lowest survival, with the majority of deaths occurring in the first year of follow-up (Figure 5B). The 75th percentile of survival in the Cluster 4 group was 90 days from the start of follow-up (95% CI: 59 to ∞ days). Cluster 3 was characterized by a high mortality rate within 1 year, but the three-year survival rate in this group was the highest. The 75th percentile of survival in the Cluster 3 group was 180 days from the start of follow-up (95% CI: 59 to ∞ days). Cluster 1 and 2 showed a more stable course of the disease and a higher survival rate than cluster 4. For the first cluster, the 75th percentile of survival was 757 days from the start of follow-up (95% CI: 302 – 1040 days), for cluster 2 it was 912 days from the start of follow-up (95% CI: 151 – ∞ days) (Figure 5B). However, the differences in overall survival between clusters as assessed by the likelihood ratio test were not statistically significant (p = 0.082075). To assess the relationship of clusters with overall survival, the Cox regression method was used.

The all-cause risk ratio for cluster 4 was significantly higher compared to other clusters (HR 2.586; 1.047 – 6.386) p = 0.041. In the presence of metabolomic cluster 4, the risk of mortality increased by 2,57 times. Cluster 3 was also characterized by a high risk of mortality (HR 1.995; CI 0.986 – 4.036) p= 0.04. (Table S10). With cluster 3, the risk of death increased by a factor of 2,119. Age was also a significant factor, increasing the risks by 1,06 times.

The analysis showed a significant increase in the risk of all-cause mortality for clusters 3 and 4. Patients belonging to clusters 3 and 4 have the worst prognosis. Thus, clustering based on metabolomic profiling makes it possible to predict the course of HF.

To determine the prognostic significance of a combination of factors: alternation of LV EF and metabolomic clusters, a model was built that included three variants of the EF trajectory during follow-up, sex, age, and metabolomic clusters. Using the Cox regression method, the assessment of the relationship between metabolomic clusters and EF alternation with overall survival made it possible to construct the following model of proportional risks (Table S11).

The factors that significantly influenced the prognosis in the model were age, decrease in EF, and the presence of cluster 3. A downward change in the trajectory of EF is associated with a threefold increase in the risk of death from all causes. Cluster 3 was associated with a 2,88-fold increase in all-cause mortality (Figure 6).

**Figure 6.**
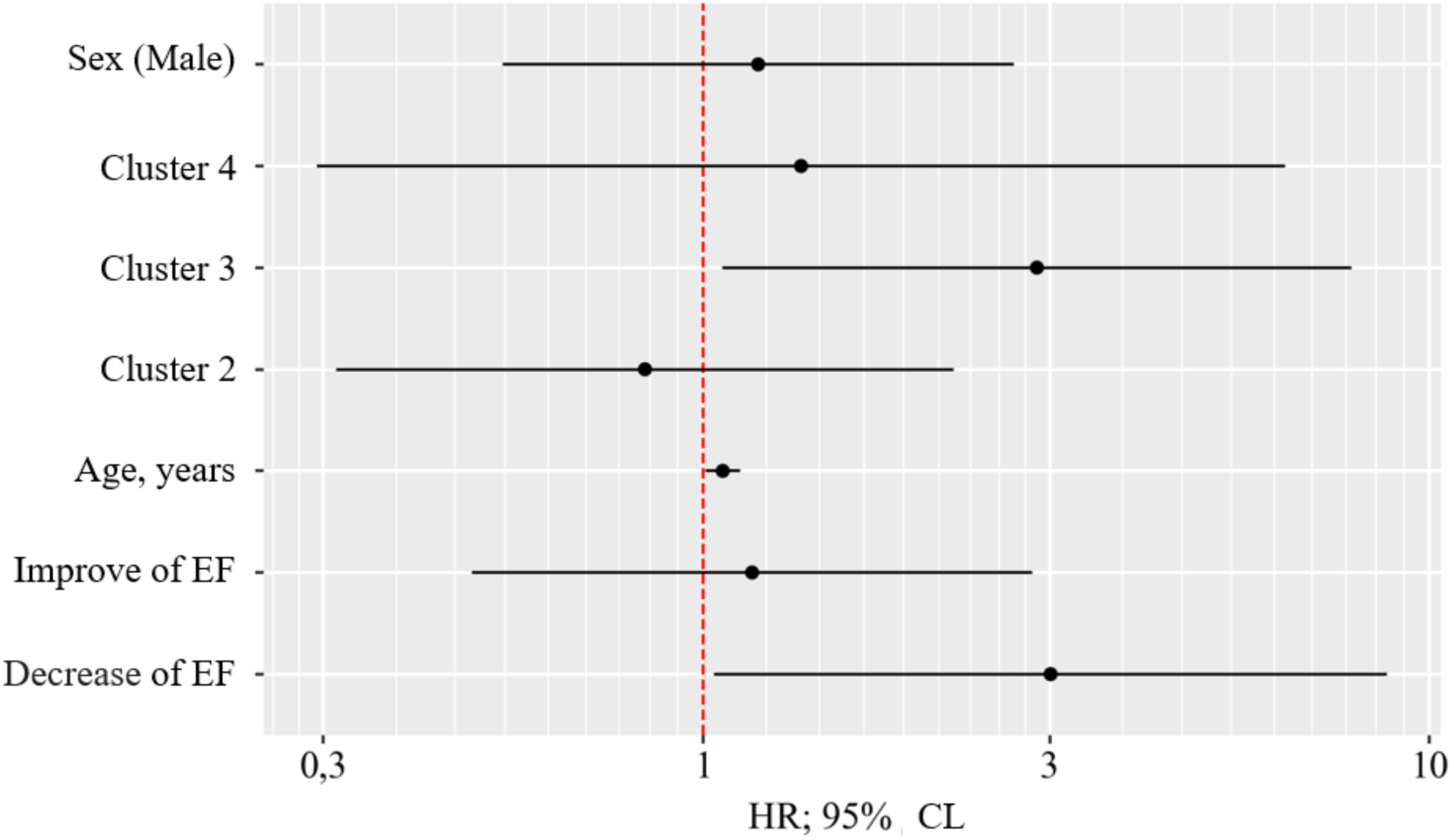
Hazard ratio estimates with 95% CI for the risk factors of all cause death. EF - ejection fraction; male - male sex.

## Discussion

To our knowledge, this is the first study to show a difference of metabolomic profile in the same grope of patients with HF, who was reclassified by few principles. Metabolomic profiling in vary stages of HF showed significant differences. The gradual decrease in nitric oxide (NO) formation (decrease in arginine and its metabolites), activation of inflammation (increase in metabolites of tryptophan and methionine catabolism), increase in oxidative stress (taurine, methionine sulfoxide, citrulline, ornithine) and a shift in energy metabolism towards a decrease in fatty acid metabolism (increase in acylcarnitine) and an increase in anaplerosis (changes in metabolites of the glutamine-ornithine-proline cycle) and glycolysis (decrease in taurine, alanine) was found during the progression of HF. The construction of a ML model made it possible to predict with great accuracy the presence of stage B or C HF. This is an extremely important observation, since the differential diagnosis of these two stages causes particular difficulties due to the no specificity of symptoms and the need to evaluate a large number of parameters and biomarkers. It is known that biomarkers, in particular NT-proBNP, can detect deterioration of the condition even before the onset of symptoms and reclassify HF from stage A to stage B.^7^ Our data demonstrate this possibility to use metabolomic profiling and ML for diagnostic of pre-HF and HF stage C. When dividing patients according to the classification based on LVEF, the results showed a significant difference in the metabolic profile of patients with HFpEF and those with LVEF <50%. At the same time, the model demonstrated low accuracy in dividing patients into groups HFpEF, HFmrEF and HFrEF. It can be assumed that metabolic disorders in HFmrEF are more similar to disorders in HFrEF than with HFpEF.

### Metabolomic profiling of HF stages

It was found that the metabolomic profile of patients with HF stage A was significantly different from the profile of patients with HF stage B. Patients with stage B were characterized by: decreased levels of neurotransmitters; an increased level of asymmetric dimethylarginine and symmetric dimethylarginine, associated with the presence of atherosclerosis and an increased levels of metabolites of the kynurenine pathway of tryptophan catabolism associated with the activation of pro-inflammatory cytokines, a decrease in metabolites of the serotonin pathway of tryptophan catabolism; increase in short and long-chain acylcarnitine’s and finally increase in trimethylamine N-oxide.^8,9^ Comparing analyses of the stage B and stage C groups, shown that symptomatic HF is characterized by a different pattern of metabolomic profile: a decrease in metabolites involved in the formation of NO. The NO–soluble guanylate cyclase (sGC)-cyclic guanosine monophosphate (cGMP) signaling pathway regulates the bioenergetics of cells and is one of the key mechanisms for maintaining homeostasis of tissues and organs.^10^ Deficiency of adenosine triphosphate (ATP), which serves as a source and energy substrate for the synthesis of nucleotides - adenosine monophosphate and guanosine monophosphate (GMP), leads to a decrease in the formation of cGMP from GMP - an activator of protein kinase G. Stage D HF compared to stage C is characterized by a significant increase in long-chain acylcarnitine’s and activation of tryptophan catabolism along the kynurenine pathway, which is probably explained by the involvement of many organs and systems in the pathological process as a result of congestion, including the kidneys and liver. The data obtained in this study was similar with the data of Mei-Ling Cheng et al., who evaluated the metabolomic profiles of patients with HF at different stages.^11^ The authors demonstrated significant differences in the concentration of metabolites in blood plasma, such as histidine, phenylalanine, ornithine, spermine, spermidine, phosphatidylcholines, and taurine, among patients at different stages of HF. As in our study, patients with the HF stage D were characterized by a violation of the glutamate-ornithine-proline cycle and a decrease in the level of histidine, arginine, and glutamine, that conferred the shift in the balance towards glycolysis. Histidine, which can be converted to glutamate, enters the Krebs cycle as alpha-ketoglutarate, which supplies ornithine.^12^ Glucose can be converted to phosphoribosyl pyrophosphate, which is essential for the biosynthesis of histidine. Excessive consumption of glucose as an energy source may impair the ability of cells to replenish histidine deficiency.^11^

Since the main goal of the work was to assess the possibility of using metabolomic profiling for the diagnosis of HF, ML methods were used as the most promising method for assessing metabolomic data and classification models of pairwise stage difference were built.^13^ As a result, it was found that the above technique makes it possible to verify stage B HF with high diagnostic accuracy both when compared with the stage A and when compared with stage C. This result is extremely important, since from a clinical point of view, the diagnosis of “pre-HF” causes great difficulties. According to the recommendations of the ACC/AHA 2022, to diagnose stage B, it is necessary to estimate many parameters: morphological (LAVI, LVMI, RWT, LV wall thickness); functional - LVEF LV global longitudinal strain, average E/eʹ, TV, PASP; biomarkers - BNP, NT-proBNP. This approach is extremely inconvenient, time-consuming and cannot be widely used in clinical practice. a simple promising way to diagnose stage B.^14^ Additional studies on a larger sample of patients are needed to assess the sensitivity and specificity of the approach proposed in our research. The low diagnostic value of the ML technique for differentiating patients from stage C and D indicates that there are no significant differences in the metabolomic profile in these patients. From a clinical point of view, additional markers for the diagnosis of stage D are not of great importance, since the clinical picture allows to differentiate stages C and D and additional treatment options will be determined primarily by the patient’s clinical status.

### Metabolomic profiling of HF phenotypes by EF

Currently, the classification method proposed by the European Society of Cardiology in 2016 and based on the identification of 3 phenotypes of HF by the level of LVEF is widely used in clinical practice.^15^ The basis for this division was an attempt to stimulate research on the characteristics, pathophysiology, and treatment approaches of patients with LVEF 40-49%.^16^ However, subsequent years have shown that HFmrEF bears features of both HFpEF and HFrEF.^16^ The randomized clinical trials conducted did not provide convincing data on the efficacy of individual classes of drugs for this group.^17–20^ Thus, the clinical guidelines for the treatment of a cohort of patients with HFmrEF suggest a management approach similar to that for HFrEF, but with a lower level of evidence.^2,3^ At the same time, it remains unclear to which cohort patients with HFmrEF should be assigned and to what extent such reclassification is pathogenetically justified.

In our own study, it was shown that the use of ML based on metabolomic profiling has low efficacy in classifying the patients by 3 EF phenotypes. This observation led to an attempt to regroup patients in two ways, assigning patients with HFmrEF to either the HFpEF group or the HFrEF group. As a result, classification models were built for two options for comparison:

1. HFpEF vs. HF and EF < 50%
2. HFrEF vs. HF and EF > 40%

It was found that metabolomic profiling makes it possible to differentiate HFpEF and HF and EF<50% with high accuracy, unlike comparison HFrEF and HF with>40%. Our data are fully consistent with the study by Wynn G. Hunter et al, which compared the metabolomic profiles of patients with HF and EF >45% and patients with systolic dysfunction (EF <45%) and showed that long-chain AC levels are elevated in patients with HFrEF compared to patients with HFpEF and significantly differ from the AC levels of healthy volunteers.^21^ In this study, significant differences were found in the concentrations of lysine, leucine, and glutamine cycle metabolites between groups amino acid elevations were significant in the group of patients with HFpEF, which is also consistent with the data of other researchers.^8,22^ Although the original study did not show the effectiveness of metabolomic profiling for differentiating patients by the three phenotypes of HF based on EF, a study by colleagues from China revealed significant differences in the metabolomic profiles of patients with HFpEF, HFmrEF, and HFrEF.^23^

Data of our study reflect a shift in metabolism towards the separation of glycolysis and tricarboxylic acid, which was demonstrated in experimental studies.^24,25^ Thus, glutamate levels were significantly increased in patients with HFpEF compared to HF with EF < 50% and in healthy volunteers. At the same time, the level of glutamate-glutamine substrate was reduced in all patients with HF compared to healthy volunteers.^26^ The accumulation of glutamate indicates a decrease in anaplerotic reactions necessary for the work of the CTA.^27^ HFpEF group was characterized by an increase in norepinephrine levels compared to the group with systolic dysfunction. It has been proven that oxidative stress is directly related to increased simpatico nervous system tone. Chronic stimulation of β-adrenergic receptors was directly related to the production of mitochondrial ROS through second-messenger signaling mediated by adrenergic receptors.^28^ Chronic sympathetic activation results in ROS-mediated initiation of mitochondrial-dependent cell death cascades.^29^ Therefore, an increase in norepinephrine levels in patients with HFpEF suggests that simpatico-adrenal system is activated and oxidative stress is exacerbated. The current findings could be interpreted in the context that the disorders of energy metabolism in HFmrEF are more similar to those in HFrEF, and the alternation of metabolism in HFpEF has its own distinctive features. Our results are confirmed by the observations of other researchers who studied the metabolomic profile of patients with HFpEF. Consequently, the data obtained cast even greater doubt on the isolation of the HFmrEF phenotype, and testify in favor of revising the classification with the allocation of two phylogroups HFpEF and HFrEF (FV<50%). Moreover, the addition of the above-described metabolites to the ML classification model made it possible to achieve a high accuracy of verification of HFpEF in comparison with HF with EF<50%.

### Metabolomic profiling for clustering HF

Using targeted, quantitative metabolomic profiling clustering of patients with symptomatic HF was carried out and 4 metabolomic clusters were identified. The construction of a ML classification model confirmed the high accuracy of the methodology for separating groups.

Previous metabolomic profiling studies confirm the association of predominantly long-chain acylcarnitines with CVDs and demonstrate their effect on the prognosis.^30–32^ For example, a systematic review published in 2017 showed that individuals with higher levels of short-, medium-, and long-chain acylcarnitines had an increased risk of CVD.^33^ This is apparently due to the importance of AC in energy metabolism, namely in the oxidation of FA, which provides up to 95% of ATP production by the heart.^34^ With a shift in the balance and a slowdown in the FA oxidation, there is an accumulation of incompletely oxidized fatty acids in the mitochondria, which contributes to lipotoxicity and intracellular acidosis, which increases the inhibition of energy production in the cardiomyocyte (CMC) and exacerbates oxidative stress, which leads to the degradation of organelles, to CMC apoptosis and ultimately to the death of a viable myocardium.^35–37^ In turn, several studies have shown a relationship between AC and HF and systolic dysfunction.^38,39^ In experimental work on mouse models, the authors describe these changes in the framework of desynchronization of glucose and FA oxidation in the CMC of hypertrophic myocardium with inhibition of mitochondrial enzyme carnitine palmitoyltransferase I/II receptor and peroxisome proliferator activated receptor.^37^

Tryptophan metabolism plays a key role in controlling hyperinflammation and inducing long-term immune tolerance. These effects are due to the ability of indolеamine 2,3- dioxygenase (IDO) to alter the local and systemic balance of kynurenin and tryptophan. This balance has a direct effect on metabolic and immune signaling pathways that regulate the anti-inflammatory response in cells that have IDO activity, such as antigen-presenting cells and epithelial cells neighboring cells, for example, T-lymphocytes, creating a local (and sometimes systemic) environment with an increased content of kynurenine and a reduced level of tryptophan.^40^ A recent study of a comprehensive analysis of tryptophan metabolism in 13 different chronic inflammatory diseases found that impaired tryptophan metabolism by kynurenine pathway (KP) is a unifying feature of chronic inflammatory diseases.^41^ There is increasing evidence linking tryptophan catabolism to the development of HF, largely due to the contribution of inflammation to the pathogenesis of cardiac remodeling and microvascular dysfunction.^42–44^ Comparing the prognostic value of NT-proBN and kynurenine in relation to the development of severe systolic dysfunction, KP demonstrated higher accuracy.^45^

### Predictors of all cause death

The disappointing epidemiological data regarding the mortality of patients with HF determined the reason to search for predictors of the risk of an unfavorable prognosis. Reduced LVEF (<40%) is a proven factor in an unfavorable prognosis. However, optimal medical treatment of HF can significantly reduce LV volumes and improve myocardial contractility, and, therefore, lead to complete normalization of LVEF (i.e., >50%) or partial recovery of LVEF (41% to 49%). ^46^ According to a systematic review and meta-analysis, 22.4% of patients with HFrEF recovered LV during treatment. HF with improved EF is associated with a 56% reduction in mortality and a 60% decrease in hospitalization rate compared to patients with HFrEF.^47^ Interestingly, when comparing the trajectories of changes in LVEF over time, it was found that patients with HF and improved EF (HFiEF) have a better prognosis even compared to patients with initially preserved EF. Patients with baseline HFpEF but who have developed a decrease in EF have the worst prognosis. The results of our research show a similar trend. Improvement in EF during therapy was noted in 35.5%. Survival analysis showed that the risk of mortality in the group of patients with a decrease in EF by more than 5%, regardless of the baseline level, was 56% with early mortality in the first year of follow-up. The three-year risk of death in all patients with HF was 41.9%. At the same time, patients with HFiEF had the highest survival rate at 3 years and amounted to 69.3%, but subsequently decreased sharply. In the present study, a significant increase in mortality in the HFiEF group after 1000 days of follow-up should be treated with caution, due to the small number of study participants, which could affect the results. Thus, our data are consistent with the results of other researchers and confirm the importance of the change the trajectory of EF in the direction of decrease, as a factor in predicting adverse clinical outcomes.

An analysis of survival between patients from different metabolomic clusters showed that clusters 3 and 4 were associated with the lowest survival, and were associated with a 2.5- fold increase in mortality risk. The most favorable prognosis was typical for patients belonging to clusters 1 and 2. Cluster 3 requires special attention, since the risk of death is highest in the first year of follow-up, which makes us think about the intensification of therapy and a possible revision of the management strategy in this group. It is important to note that metabolomic clusters did not differ in the trajectory of EF changes. The combination of the assessment of the trajectory of EF and the cluster made it possible to identify factors that affect the prognosis: a decrease in EF in dynamics, the presence of cluster 3, and an increase in age. To assess the sensitivity and specificity of the method, additional studies are required in a larger cohort of patients.

Thus, in order to improve the prognosis of patients, it is necessary to take into account such factors as a decrease in EF during patient follow-up by 5% and the presence of the cluster 3 metabolomic phenotype and think about a possible revision of the therapeutic strategy or its intensification in this groups of patients.

### Clinical perspectives

To our knowledge, this is the first study to explore the potential of ML based on metabolomic profiling technics to analyze HF across a full spectrum of stages and EF phenotypes and the first to apply clustering to HF using metabolomic profiling. Modern diagnostic methods, combining mass spectrometry in combination with ML, new avenues for precise identification of HF subtypes, the discovery of unique phenotypic characteristics, more accurate prediction of disease prognosis and complication risks, and automated analysis of large datasets.

In this study, the use of metabolomic profiling for HF diagnosis was evaluated in line with existing classification systems. Metabolomic analysis confirmed the utility of ML classification models in differentiating HF stages and EF-based phenotypes. These AI-driven supervised ML classification tools provide novel diagnostic options, independent of traditional classifications. Additionally, metabolomic data support the phenotypic similarity between HFmrEF and HFpEF, reinforcing the argument for a revised classification system that divides patients into two primary groups, HFpEF and HFrEF. However, further research is needed to establish the EF threshold to best differentiate these groups. Notably, changes in EF trajectory during treatment remain a key predictor of prognosis and should be integrated into patient management strategies.

Clustering data are particularly valuable as they reveal new HF phenotypes with distinct pathophysiological mechanisms, offering a foundation for developing targeted treatment approaches for these patient subgroups.

### Limitations

The main limitation of this study is the relatively small sample size, which may affect the accuracy of the ML classification model. Repeated echocardiography was not performed for all study participants but only for those who regularly visited the center during the observation period. Consequently, data on changes in ejection fraction over time may be incomplete, limiting the assessment of temporal dynamics.

### Conclusions

The plasma metabolome reflects the primary metabolic changes occurring in the body and serves as an imprint of the phenotype. The metabolic similarities observed between the HFrEF and HFmrEF phenotypes underscore the limitations of the current classification and suggest the need to revise it, potentially combining patients into two main groups. At the same time, the high diagnostic accuracy for detecting HF stages B and C using ML based on metabolomic profiling supports the effectiveness of this approach for refining HF stage classification and diagnosis.

HF phenotyping through hierarchical clustering based on metabolomic profiles demonstrated high model accuracy. Further studies are needed to clarify the role of metabolomic profiling as a diagnostic tool in HF and to identify patients who may benefit from metabolism-targeted therapies. The findings confirm that metabolomic profiling methods present a promising alternative for patient stratification and disease classification.

## Acknowledgments

The authors express great gratitude to the doctors of the Department of Cardiology No. 1 of the University Clinical Hospital No. 1 of Sechenov University for their assistance in recruiting and clinical supervision of patients. The graphical abstract was created with www.BioRender.com.

## Sources of Funding

The study was conducted with financial support of the Innovative Science School at Sechenov University within the framework of the *Priority 2030* program.

## Conflict of interest

none declared.

## Disclosures

None.

## Supplemental Material

Methods.

Figures S1.

Tables S1–S11.

## Nonstandard Abbreviations and Acronyms

AC: acylcarnitines
ACEi: angiotensin-converting enzyme inhibitor
ARB: angiotensin receptor blocker
ARNI: angiotensin receptor-neprilysin inhibitor
ATP: adenosine triphosphate
AUC ROC: area under the receiver operating characteristics curve
BMI: body mass index
BNP: brain natriuretic peptide
CAD: coronary artery disease
CABG: coronary artery bypass grafting
CCB: calcium channel blocker
CMC: cardiomyocyte
cGMP: cyclic guanosine monophosphate
CVD: cardiovascular diseases
CRP: C-reactive protein
e’: early diastolic velocity of movement of the fibrous ring of the mitral valve
EF: ejection fraction
eGFR: estimated glomerular filtration rate
FA: fatty acids
GDMT: guideline-directed medical therapy
HCA: hierarchal cluster analysis
HDL: high-density lipoprotein
HF: heart failure
HFiEF: heart failure and improved ejection fraction
HFmrEF: heart failure with mid-range ejection fraction
HFpEF: heart failure with preserved ejection fraction
HFrEF: heart failure with reduced ejection fraction
IDO: indoleamine 2,3-dioxygenase
KP: kynurenine pathway
LA: left atrium
LAVI: left atrium volume index
LDL: low-density lipoprotein
LC-MS/MS: liquid chromatography mass spectrometry
LV: left ventricular
LVEF: left ventricular ejection fraction
LVMI: left ventricular mass index
MI: myocardial infarction
ML: machine learning
MP: metabolomic profiling
MRA: mineralocorticoid receptor antagonist
NO: nitric oxide
NT-proBNP: N-terminal pro-B-type natriuretic peptide
PCA: principal components analysis
PCI: percutaneous coronary intervention
QC: quality control
RWT: relative wall thicknes
sGC: soluble guanylate cyclase
SGLT2i: sodium/glucose cotransporter-2 inhibitors
SPAP: systolic pulmonary artery pressure
TR: tricuspid regurgitation
TIA: transient ischemic attack
WBC: white blood counts

## Notes

### Competing Interest Statement

The authors have declared no competing interest.

